# Unexpected sound omissions are signaled in human posterior superior temporal gyrus: an intracranial study

**DOI:** 10.1101/733212

**Authors:** Yvonne M. Fonken, Arjun Mukerji, Richard Jimenez, Jack Lin, Peter Brunner, Gerwin Schalk, Robert T. Knight

## Abstract

Context modulates sensory neural activations enhancing perceptual and behavioral performance and reducing prediction errors. However, the mechanism of when and where these high-level expectations act on sensory processing is unclear. Here, we isolate the effect of expectation absent any auditory evoked activity by assessing the response to omitted expected sounds. Electrophysiological signals were recorded directly from the superior temporal gyrus (STG) and superior temporal sulcus (STS) in patients with medically refractory epilepsy. Subjects listened to a predictable sequence of syllables, with some infrequently omitted. We found a high frequency band (HFB, 70-150Hz) response to omissions, which overlap with a posterior subset of auditory active electrodes. This response is distinct from omission activations observed in non-auditory selective sites in STG. Heard syllables could be classified reliably from STG, but not the identity of the omitted stimulus. Both omission- and target detection activations were also observed in prefrontal cortex.

We propose that the posterior STG and STS are central for implementing predictions in the auditory environment. HFB omission activations in this region appear to index mismatch-signaling or salience detection processes.

## Introduction

### Expectations influence sensory processing

The notion that the brain uses prior knowledge to make predictions about incoming sensory input has gained considerable traction (Friston 2009; Friston 2010; Arnal and Giraud 2012). The idea is that the brain does not process incoming sensory signals in a purely feedforward manner as previously believed (Serre et al. 2007), but implements cortico-cortical feedback that influences sensory processing in a top-down, hierarchical manner (Rao and Ballard 1999; Lee and Mumford 2003). The advantage of a prediction strategy is improved perception and behavior (Anllo-Vento 1995; Mangun 1995). On a behavioral level, prior knowledge enhances intelligibility of noisy speech. The underlying mechanism for this effect includes rapid expectation-dependent changes in auditory perceptive field responses (Holdgraf et al. 2016). Evidence of expectations influencing early sensory processing have also been shown in vision as reductions of the V1 BOLD response to expected gratings (Alink et al. 2010) and expected tones reduce the auditory N100 amplitude in MEG recordings (Todorovic et al. 2011).

### Prediction and the brain

The idea of the brain as a prediction machine was first proposed by Helmholtz, and has often been described in a hierarchical Bayesian framework in computational models (Rao and Ballard 1999; Lee and Mumford 2003; Friston 2005; Olshausen 2014). This view describes how predictions and expectations based on prior knowledge aid perception and action. It is suggested that higher order cortical regions communicate predictions to lower-order regions hierarchically through a multitude of recurrent connections (Clark 2013). One specific theory of the predictive brain is termed predictive coding (Rao and Ballard 1999; Friston 2005). In the predictive coding framework, local neuronal populations compute errors based on top-down predictions, and these prediction errors are propagated up the hierarchy (bottom-up) to influence subsequent behavior (Rao and Ballard 1999; Friston 2005). According to Friston & Kiebel (2009), the brain operates to ‘explain-away’ signals from lower levels of processing, providing an account for reduced responses to expected stimuli. However, a discrepancy in the literature arises when on the one hand predictions are proposed to reduce neural responses lower in the sensory hierarchy (Friston 2010), yet predictable stimuli are more easily decoded from V1 voxels despite smaller BOLD responses (Kok et al. 2012). This suggests that predictions may not simply reduce neural activity in sensory processing areas, but perhaps facilitate processing the expected stimulus by enhancing stimulus-specific information (Kok et al. 2012).

### Investigating auditory context processing through omissions

Studies investigating predictions often manipulate stimulus predictability or embed stimuli in noise, and the resultant auditory activity is a confluence of bottom-up sensory processing, and expectation modulations. Here, we aimed to isolate expectation effects on auditory cortex by examining the neural signals to omissions of expected sounds. Omissions of expected sounds have been shown to elicit ERP responses in EEG ~100ms after the expected sound onset, which are generated in auditory cortices (Sanmiguel et al. 2013; Bendixen et al. 2014). In the visual domain, omission signals in V1 have been shown to contain stimulus-specific information since the omitted stimulus can be decoded from V1 voxels using fMRI. This has been interpreted as an activation of a stimulus template (Kok et al. 2014). In auditory cortex, omitted speech sounds embedded in words can also be recovered from HFB activity in STG. The omitted word sections were replaced by noise, yet HFB reconstructions of the omitted section matched the perceptual experience of the subject (Leonard et al, 2016). Evidence for higher-order information influencing human auditory STG HFB response patterns has also been shown in a study using noisy stimuli that become intelligible in the presence of prior knowledge of what is presented (Holdgraf et al. 2016). HFB activity has been shown to drive the fMRI BOLD response and correlates with neural firing, providing a link between different methods (Niessing et al. 2005; Ray and Maunsell 2011). Here, we utilized the high spatial and temporal resolution of ECoG to: 1) isolate prediction-related HFB activations to auditory omissions in human auditory cortex; 2) define the spatiotemporal dynamics of these activations; and 3) determine whether this HFB activation carries stimulus-specific information.

## Materials and Methods

### Participants and experimental setup

A total of 6 subjects (1 female, mean age 43, range between 31 and 69) participated in the current study (see Supplementary Table 1 for further demographic information). All subjects had extensive coverage of the lateral STG, except IR01, who only had depth electrodes in STS. An overview of electrode coverage for AB01-AB05 can be seen in Supplementary Figure 1. Subjects were recruited from a patient group with medically refractory epilepsy undergoing neurosurgical treatment, and had subdural electrodes implanted for clinical purposes. These patients were tested during clinical monitoring in their hospital bed, and typically remained implanted for a duration of 4-10 days. All patients gave their informed consent according to the Declaration of Helsinki, and an additional verbal consent was given prior to each testing session. Patients were recruited from different sites, including Albany Medical Center (AB, n=5) and UC Irvine (IR, n=1). Institutional Review Boards from each individual site and UC Berkeley approved the experimental procedures.

Electrophysiological signals were recorded using either a g.tec g.HIamp system and digitally sampled at 9.6 kHz (AB) or a Nihon Kohden system with a 128-channel JE-120A amplifier at a sampling rate of 5 or 10kHz (IR). Electrodes were made of platinum-iridium and spaced 3-10mm, and were manufactured by AdTech or PMT. Depth electrodes were spaced 5mm apart, and were manufactured by AdTech. Reconstructions of electrode placement were made using pre-operative T1 structural MRI scans and post-operative CT scans using Bioimagesuite (Irvine) or the matlab toolbox SPM8 (Albany). Average brain projections were done using the Fieldtrip toolbox (Oostenveld et al. 2011). Timing of stimuli was recorded in analog channels by splitting the speaker signal from the experimental computer to the recording system. Responses in the form of finger taps were recorded using a microphone plugged into an analog channel (IR), or using a response button (AB). Analog channels were recorded at 5kHz or higher.

### Experimental Task

To enhance stimulus predictability, we played a repetition of the pattern ‘La-La-Ba La-La-Ga’ using syllable stimuli created and shared by the Shannon lab at USC. We chose to use syllables as stimuli to ensure robust auditory activations in the STG. We chose ‘Ba’ and ‘Ga’ since these syllables have been previously shown to be decodable from this region (Chang et al. 2010). We used a third syllable ‘La’ to set up a temporal expectation of the ‘Ba’ or ‘Ga’ to be played. To ensure that the subject was attentive to the sounds, the subject was instructed to respond to a ‘Ta’, which we randomly introduced in place of the ‘Ba’ or ‘Ga’ as a target stimulus 5% of the time. Since the task is very repetitive, we chose a target stimulus close to the other stimuli to prevent the subject from ignoring the stimuli and simply relying on bottom-up salience of a target stimulus. Finally, the relevant task manipulation was the omission of either ‘Ba’ or ‘Ga’. The syllables lasted 410ms each, and the ISI within a ‘La-La-Ba’ triplet was fixed to 200ms, whereas the ISI between triplets was 400ms (Irvine) or 200ms (Albany). We recorded between 3 and 6 blocks in each subject, with each block including 16 omission trials, 8 target trials, and 68 ‘Ba’ and ‘Ga’ presentations respectively, and lasted about 4 minutes. The experiment was coded using E-Prime 2.0 software (Psychology Software Tools, Pittsburgh, PA) (Irvine), or BCI2000 (Schalk et al. 2004) (Albany).

### Preprocessing

The data was analyzed using custom-written scripts and MNE, scipy, numpy, and pandas packages in Python (Gramfort et al. 2013). All data was down-sampled to 1kHz (IR) or 1.2kHz (AB) after a 500Hz low-pass filter, corrected for DC shifts, band-pass filtered between 0.5Hz and 220Hz, and notch-filtered to remove line-noise at 60Hz, 120Hz and 180Hz, using a FIR filter from the MNE Python toolbox (Gramfort et al. 2013). AB data was filtered at 1Hz to 190Hz due to intermittent high-frequency machine noise above this frequency. All channels were re-referenced to a common average (surface electrodes), or were bipolar referenced (depth electrodes). Timing of stimuli and responses was extracted from the analog channels. Omission onsets were calculated by subtracting the onsets of the previous two stimuli (‘La-La’), and added to the onset of last stimulus (second ‘La’).

### Electrophysiological analysis

Spectral power was calculated using Hilbert transform after band-pass filtering between 70-150Hz. Statistical evaluations were done on the single trial analytic amplitudes. We employed a non-parametric clustering algorithm (Oostenveld et al, 2008) on a time-window of 0-500ms comparing single trial analytic amplitudes in omission trials to a baseline of 200ms to 50ms before the stimulus was expected. The power percent change was calculated relative to an average of a baseline taken 200ms to 50ms before stimulus onset. Time-course plots were made using the Seaborn Python visualization package, with error bars signifying a 68% confidence interval bootstrapped across trials.

Analysis concerning the location of effects along the anterior-posterior axis were done by localizing and visualizing electrodes using the Fieldtrip toolbox, which is part of the pipeline for the integrated analysis of human intracranial data (Oostenveld et al. 2011; Stolk et al. 2018). Electrodes were normalized and projected onto a Freesurfer template fsaverage brain using surface-based normalization (Dale et al. 1999). Electrodes were selected based on whether significant HFB increases were observed compared to baseline (within electrode cluster permutation, 0-500ms, p<0.05). Electrodes were grouped into two categories: electrodes with significant omission and auditory HFB increases, and electrodes with significant auditory activations but no significant omission activations. This distinction then served as an independent variable in a linear-mixed effects model, with the MNI Talairach y-coordinate as the dependent variable, and subject as a random effect on the intersept to control for inter-individual differences.

For the peak latency analysis, first the average trace per electrode was filtered using a low-pass filter at 10 Hz to smooth the data. The latency of the maximum amplitude was determined with a max function from numpy. The peak latency was calculated for each individual electrode for Ba and Ga trials together as well as for omissions. Latencies were first averaged across electrodes for each subject, and subsequently averaged across subjects. In addition, latency differences were calculated at the electrode level, subtracting the Ba/Ga peak latency from the omission latency in that specific electrode. Latency difference were then averaged across electrodes for each subject, and subsequently averaged across subjects. A t-test was done on the subject averages for electrodes displaying both omission and auditory responses.

For binary classification, we used a linear support vector machine classifier from the scikit-learn Python toolbox. Prediction accuracies were obtained using a 10-fold cross-validation. The first classification approach involved an electrode selection based on how good each electrode classifies Ba vs Ga (actual sounds). The purpose of this classification approach was to test if auditory sites distinguishing between the heard sounds also distinguish which stimulus was omitted. A separate classifier for each electrode was applied on its HFB data from 0-500ms relative to stimulus onset (see Fig. 4). If an electrode had a prediction accuracy of more than 60% based on hearing the spoken sounds, it was included in omission classification. For omission classification, this selection of electrodes’ HFB data from 0-500ms relative to omission ‘onset’ was included in a single classifier. Both training and testing were done on omission trials. A second classification approach included features of omission HFB data that survived a statistical threshold compared to baseline. This specifically tested whether HFB increases to omissions carried information on which stimulus was omitted. The statistical threshold consisted of applying non-parametric permutation clustering on individual electrodes in a time-window of 0-500ms, comparing omission HFB to baseline. HFB activity for significant clusters was subsequently used as features to classify which stimulus was omitted.

## Results

### Behavioral results

Omissions were embedded in a predictable stream of syllables (‘La-La-Ba’; ‘La-La-Ga’) in a target detection task to ensure attention. Occasionally either the ‘Ba’ or ‘Ga’ syllable was omitted, or replaced by a target syllable (‘Ta’) that required a button press. The behavioral data in Supplementary Table 2 show that subjects responded to targets with average reaction times ranging between 438ms and 643ms (552ms ± 156ms). The hit rate across subjects was 90%, with the lowest individual hit rate at 81%. Subject AB03 had a larger number of false alarms (43 trials) compared to the other subjects, i.e. button presses to ‘Ba’ or ‘Ga’.

### High Frequency Band activations in auditory regions to syllables

Local field potentials were recorded directly from human STG and STS using ECoG grids and depth electrodes in six patients undergoing clinical evaluation for medically refractory epilepsy (for electrode coverage see Supplementary Figure 1). High frequency band (70-150Hz; HFB) task activations in auditory regions for individual subjects are shown in Figures 1 and 2. Figure 1 shows two subjects with high-density grids (3mm inter-electrode spacing) covering the STG. Panels A and C (left side) show that auditory activations to syllables evoke a robust HFB activation in the STG and STS. Auditory activations typically onset <100ms, as can be seen in the time courses plotted in panels B and D (blue and red traces). These activations were significant compared to baseline with clusters in 87 electrodes for AB01, and 98 electrodes for AB02 (cluster permutation, 0-500ms, p<0.05). Additional subjects show similar patterns, as can be seen in Figure 2A and 2C. Here, AB03 showed significant HFB activations compared to baseline in 31 electrodes, AB04 in 15, AB05 in 46 electrodes (cluster permutation, 0-500ms, p<0.05, topographies in Fig. 1 and 2 show data of significant activations only). In addition, we also found robust auditory activations in the superior temporal sulcus (Figure 2, subject IR01).

**Figure 1.**
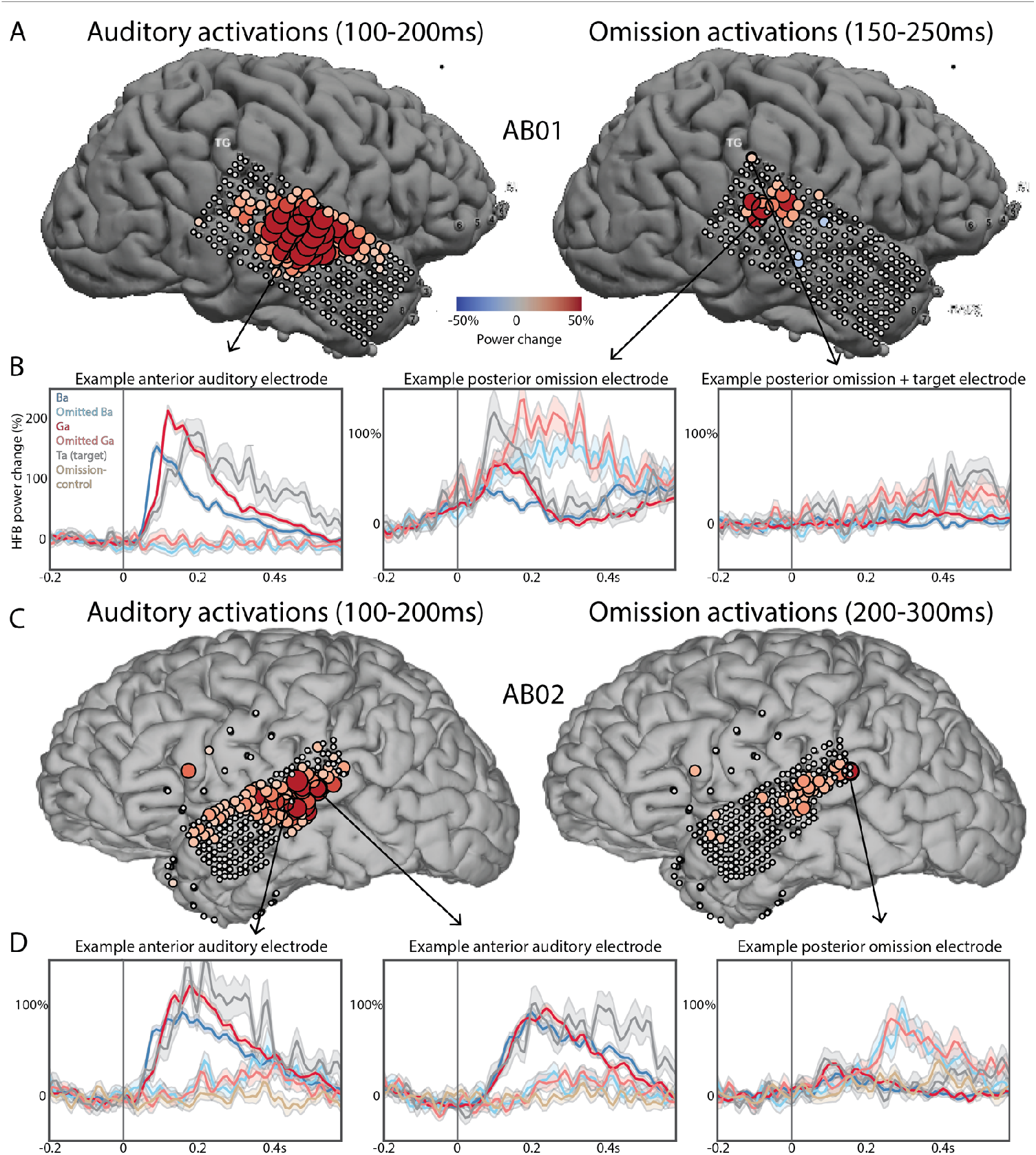
HFB activation patterns for spoken and omitted sounds for two high-density (3mm) grid subjects. A) Topography of HFB activations for subject AB01. HFB power is averaged for (omitted) ‘Ba’ and ‘Ga’ presentations over a time-window of 100-200ms (left) and 150-250ms (right). B) Three example electrodes showing: 1) auditory but no omission activations; 2) auditory and omission activations; and 3) omission and target activations but no auditory activations. Stimulus onset is at 0ms, and traces are HFB responses to ‘Ba’ (dark blue), ‘Ga’ (dark red), omitted ‘Ba’ (light blue), omitted ‘Ga’ (light red), ‘Ta’ (gray) and an omission control (tan). C) Topography of HFB activations for subject AB02. HFB power is averaged for (omitted) ‘Ba’ and ‘Ga’ presentations over a time window of 100-200ms (left) and 200-300ms (right). D) Three example electrodes show both auditory and omission activations.

**Figure 2.**
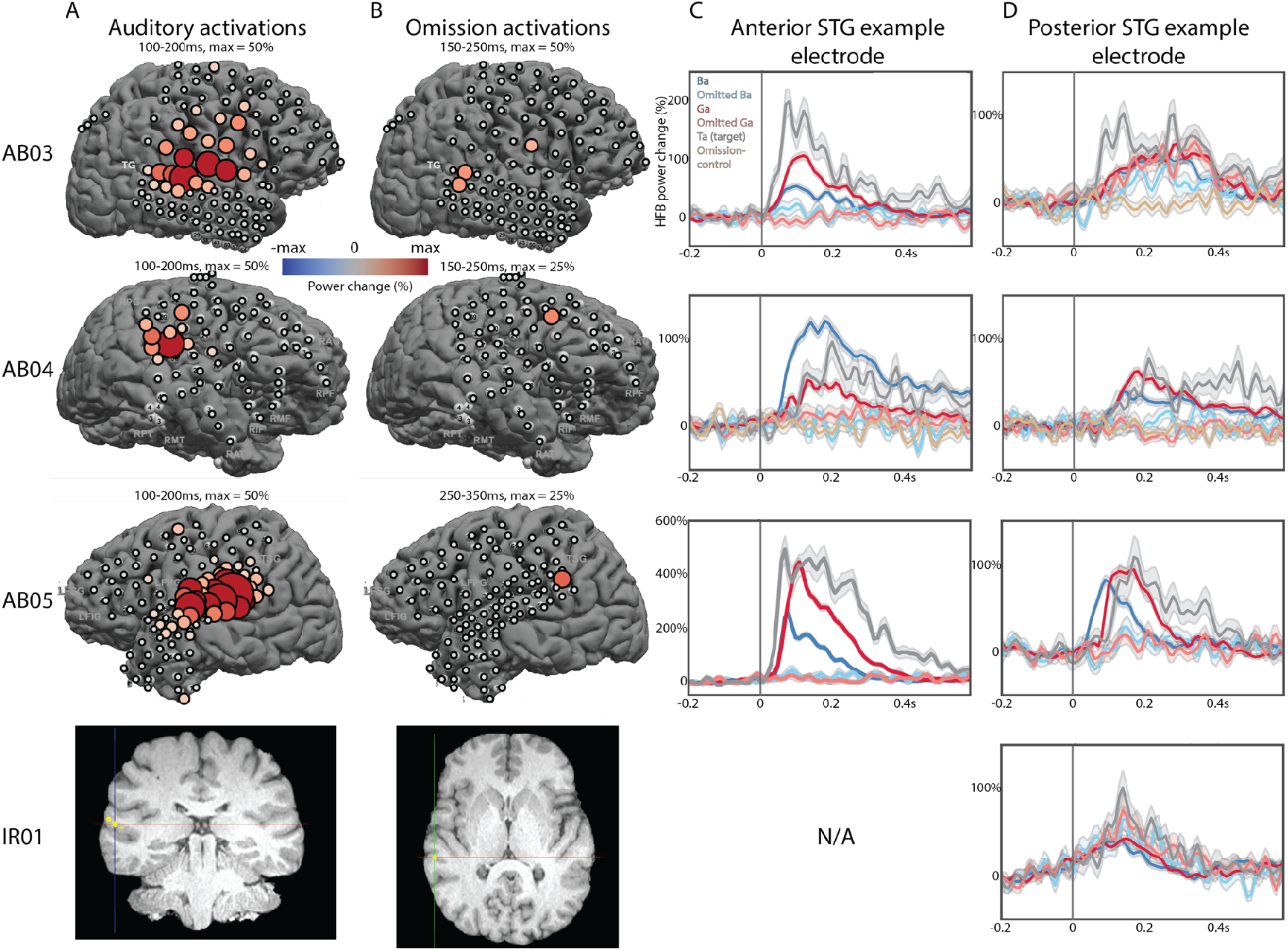
Auditory and omission HFB activations in additional subjects. A) Topography of significant auditory HFB power change. Specific time-window and scale are shown above each plot. Subject IR01 only shows the electrode location. B) Topography of significant omission HFB power change. C) Example auditory electrodes in anterior STG. Stimulus onset is at 0ms, and traces are HFB responses to ‘Ba’ (dark blue), ‘Ga’ (dark red), omitted ‘Ba’ (light blue), omitted ‘Ga’ (light red), ‘Ta’ (gray) and an omission control (tan). D) Example auditory electrodes in posterior STG (or STS for IR01).

### High Frequency Band activations to omissions in auditory regions

Responses to omissions are shown in Figure 1 and 2 (light-colored red and blue traces). HFB increases to omissions predominantly occurred in posterior subsets of auditory active electrodes (see Fig. 1A,C; right side). Example anterior electrodes show robust auditory activations, whereas omission HFB power shows no or small increases (Fig. 2C, Fig. 1B, D). In contrast, posterior auditory active electrodes show robust power increases in response to both omissions and sounds. HFB power increases were statistically tested compared to baseline (−200 to −50ms) within subjects using a cluster permutation algorithm to control for multiple comparisons (Maris and Oostenveld 2007). For omissions, significant clusters (cluster permutation, 0-500ms, p < 0.05) were found in 28 electrodes in AB01, and 37 electrodes in AB02, in a time-window of 0-500ms. In subject AB01 (Fig. 1B, second plot), the HFB power deviated from baseline at 0ms suggesting anticipatory activity to the expected time of the stimuli. However, this early HFB increase was not observed in lateral STG in other subjects.

Figure 2 also shows omission activations in subjects AB03 (3 electrodes, cluster permutation, 0-500ms, p<0.05), AB05 (5 electrodes) and IR01 (1 electrode), but no significant omission activations in auditory electrodes in AB04. Note that these omission responses are not limited to the lateral surface of the superior temporal gyrus, as subject IR01 shows a similar response in a depth electrode in STS. In some cases, the omission activation is similarly sized or larger compared to the auditory activations (Fig. 1B&D). Some electrodes in STG showing omission activations are unique in that they show both omission and target activations, but no auditory activations (Fig. 1B, third example electrode). No STG omission responses were recorded in subject AB04 who had the sparsest coverage.

To test whether auditory- and omission-active electrodes were more posterior compared to auditory-only STG electrodes, we compared MNI y coordinates of these two groups of electrodes in subjects with extensive STG coverage and including omission responses AB01, AB02, AB03, and AB05. The mean y coordinate difference between omission and auditory active electrodes compared to auditory-only electrodes is −15.6 ± 7.5 (calculated across subject averages). To test significance, a linear mixed effects model was used on all electrodes, with subject as random effect on the intercept. In this model, auditory and omission electrodes were significantly more posterior compared to auditory-only electrodes (p<0.0001, with a coefficient of −10.6 for auditory & omission relative to auditory-only electrodes). In all subjects, sites with auditory and omission responses are consistently more posterior than sites responding to auditory stimuli only, as can be seen in Figure 3 and in Supplementary Figure 3.

**Figure 3.**
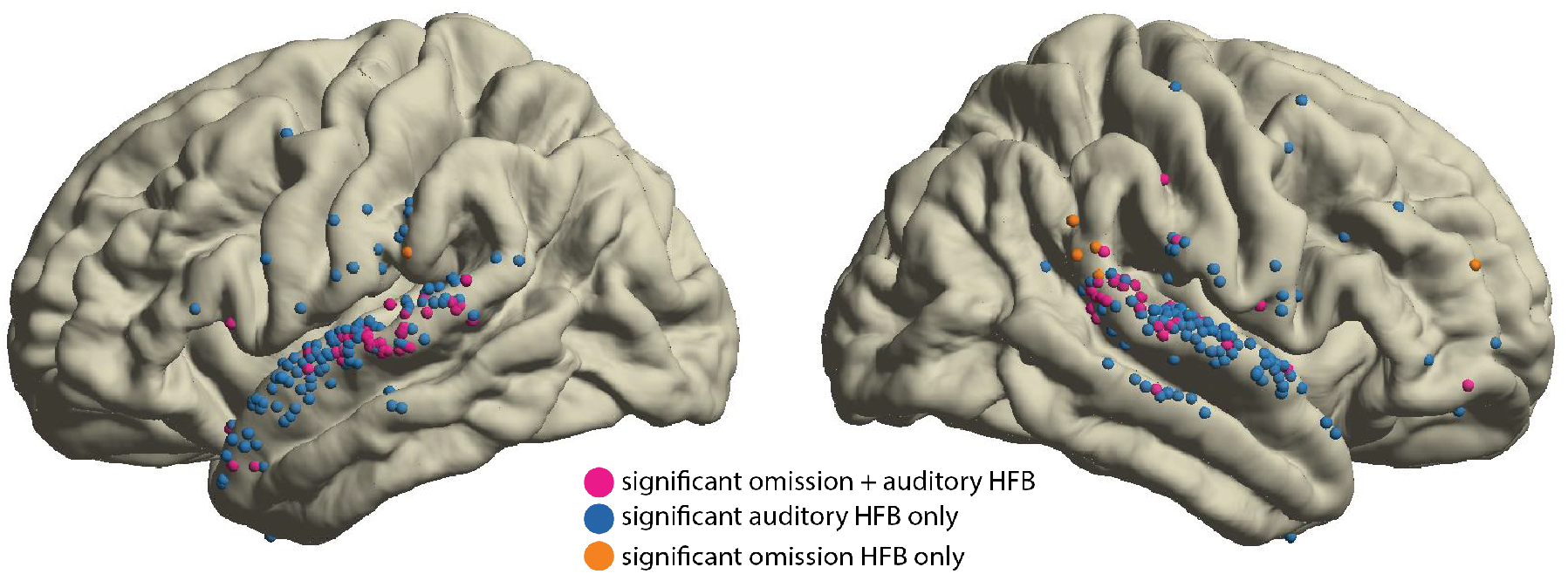
Overview of results showing omission responses in posterior auditory active sites, including electrodes from subjects AB01, AB02, AB03, AB04, AB05, projected onto the fsaverage brain from the Fieldtrip toolbox. Magenta colored electrodes are electrodes with both significant auditory and omission HFB power increase versus baseline, blue colored electrodes signify electrodes with significant auditory HFB power increases only, whereas orange colored electrodes only show significant omission HFB power increases (0-500ms, p<0.05 cluster permutation)

The omission response peaked later than auditory-evoked HFB activations. Average omission response peak latencies across subjects ranged between 238ms and 326ms, with an average of 287ms ± 40ms. Auditory responses in electrodes displaying omission responses showed peak latencies ranging between 157ms and 192ms, averaging 174ms ± 14ms. This is comparable to auditory response peak latencies in auditory electrodes that did not display omission activity, at 177ms ± 28ms. Differences in latencies between auditory and omission responses were calculated within electrodes, and subsequently averaged per subject. Across subjects, omission responses peaked between 72ms and 170ms later than auditory responses, averaging 113ms ± 41ms (p<0.02, T = 4.72, paired t-test, N=4 subjects). This difference in latencies can be observed at a single electrode level (Fig 1B, D; Fig. 2D), and is consistent across subjects (Supplementary Figure 4). Overall, omission HFB responses in STG are observed in posterior auditory-active sites, and peak ~100ms later than auditory responses.

### Decoding analysis using auditory HFB activations to sounds and omissions

Decoding analysis of spoken sounds was successful for heard syllables at the single electrode level, with prediction accuracies >80% in some electrodes (see Fig. 4). Ba versus Ga classification was successful based on significant HFB increases to syllables including all electrodes. Classification accuracy across subjects was 84.4% ± 13.6%, compared to a 99.5% confidence interval at 54.9% ± 0.3% based on a random permutation distribution (see Supplementary Table 3). To test whether auditory sites distinguishing between ‘Ba’ and ‘Ga’ can also distinguish which of these two syllables were omitted, we employed a classifier on a time-window of 0-500ms including all electrodes with prediction accuracies >60% for heard ‘Ba’ and ‘Ga’. This classification approach averaged 45.3% ± 3.7% across subjects, while the 95% confidence interval in the null distribution was at 59.7% ±0.7%. This chance level performance indicates that we were not able to determine the identity of the omitted stimuli from HFB activity in syllable-selective sites in STG.

**Figure 4.**
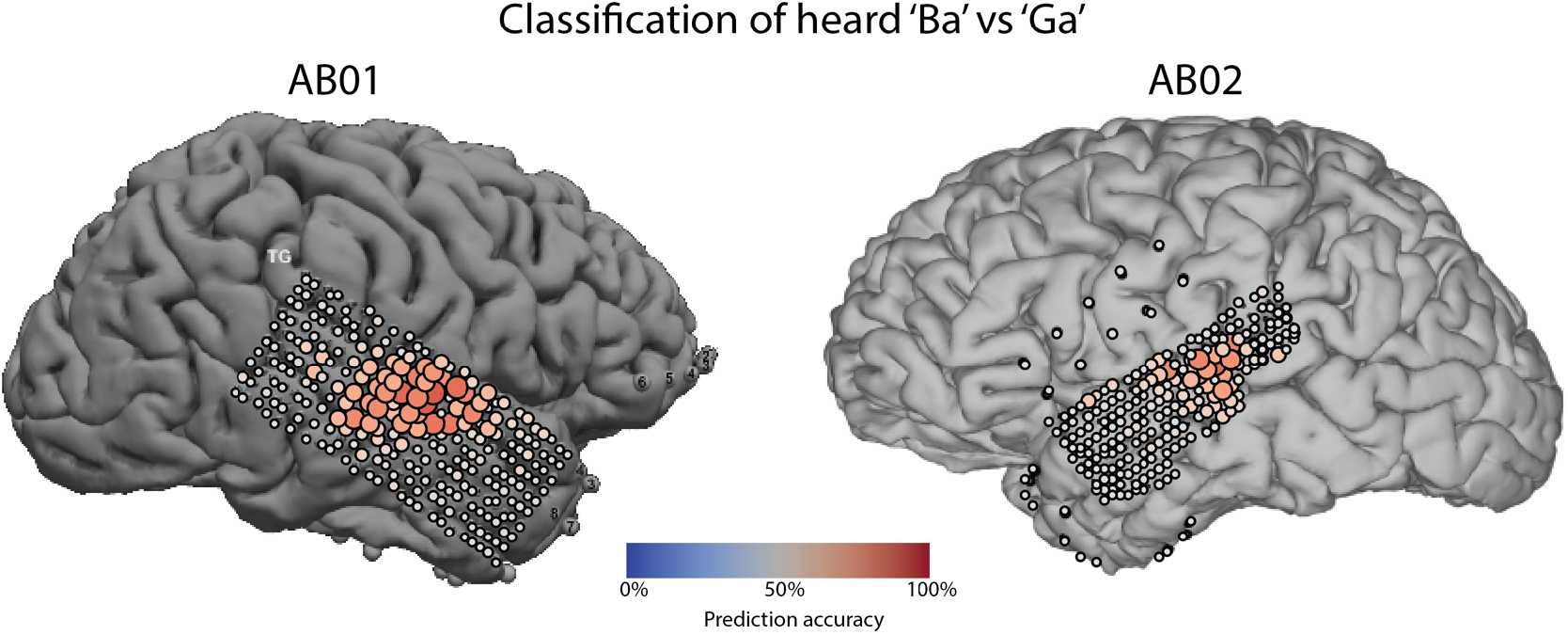
Single electrode classification prediction accuracies for classifying heard Ba vs Ga in subjects AB01 and AB02. Despite good heard Ba vs Ga classification performance as shown here, classification of which sound was omitted was not successful (data not shown).

Since omission HFB activity was observed in only a subset of auditory active sites, our electrode selection based on successful syllable decoding may not be the optimal feature space for decoding which stimulus was omitted. This is because electrodes with omission activations do not fully overlap with the electrode selection based on good ‘Ba’ vs ‘Ga’ classification. We therefore tested if HFB increases to omissions in STG specifically indexed which stimulus was omitted. Electrodes and time windows were selected based on HFB power comparisons to baseline using the cluster permutation method within electrodes in a time-window of 0-500ms. All significant clusters were subsequently included as features to classify which stimulus was omitted. This approach using multiple electrodes with only omission activity also remained at chance level for all subjects, with an average of 48.9% ± 6.9% and an average 95% confidence interval of 59.6% ±1.0%.

### HFB activity in STG to target stimuli

Target sounds violate predictions. Target syllable (‘Ta’) HFB activations were compared with HFB activations to ‘Ba’ and ‘Ga’ to investigate potential mis-prediction effects. Target sounds elicited an enhanced HFB auditory response (onset 50-150ms) in some STG electrodes for four out of five subjects with STG coverage. After the initial evoked response, HFB power remained elevated for target stimuli up to ~600ms post-stimulus (for examples see Fig 1B, D, Fig. 2C,D). In addition, target stimuli also elicited HFB activations in rare prefrontal cortex sites (see Supplementary Figure 2). To investigate whether these differences overlapped with omission activations in STG, the ratio between target stimulus and ‘Ba’/’Ga’ HFB power was compared between omission-active auditory electrodes and omission-silent auditory electrodes (data not shown). We did not identify consistent differences between omission-active versus omission-silent auditory electrodes with respect to target stimulus activity.

## Discussion

We used intracranial recordings to investigate expectation effects on auditory cortex. We isolated stimulus-independent neural prediction activity by examining omissions of expected sounds. We found a posterior subset of auditory active electrodes in the STG with robust HFB power increases to omissions (Figure 1–3). However, information on which stimulus was omitted was not encoded in these HFB power increases.

### Posterior STG activates to omissions of expected sounds

HFB power increases were elicited by omissions of expected speech sounds. These activations were observed in all subjects except AB04, which may be due to limited coverage over posterior auditory areas in STG. The most notable finding was that omission activations were observed in a posterior subset of auditory-active electrodes in the STG, which were distinct from omission activations in non-auditory responsive sites further posterior in STG (Figs. 1–3). Because we used spoken sounds as stimuli, we could uniquely map out auditory processing regions in STG that may not respond to simpler stimuli, such as tones that often are used in other mismatch studies (Edwards et al. 2005; Dürschmid et al. 2016). These results challenge current prediction error signal accounts and theories of sensory cortex dependent mechanisms of predictive coding. In these models, a prediction error is hypothesized to be produced and propagated along the hierarchy of sensory processing (Friston 2010; Bastos et al. 2012). However, our HFB omission activation was generated only in a subset of auditory responsive sites, and HFB power to omissions remained at baseline in the majority of STG electrodes that responded strongly to speech sounds.

This anterior versus posterior separation of omission activity in the STG may be related to anatomical separations of the auditory processing stream. A study by Ozker et al (2017) (Ozker et al. 2017) shows that noisy speech in the presence of contextual cues differentially affects posterior and anterior STG. HFB responses to speech in posterior STG was unaffected by added noise with context, whereas auditory activations degraded in anterior STG, suggesting an audio-visual integration role for posterior STG. In our task, contextual information arose from knowledge of the stimulus sequence structure, supporting a role of this region in contextual processing. Damage to the posterior STG and angular gyrus has been associated with a specific auditory short-term memory deficit (Markowitsch et al. 1999; Gazzaniga et al. 2002). The posterior STG may be supporting auditory short-term memory critical for recognizing patterns and signaling deviations.

An anterior-posterior division has also been reported in right STG for consonant compared to dissonant cords processing (Foo et al. 2016). This division has also been reported for spoken sentences, with the posterior STG responding mostly to onsets of sentences and emphasizing syllables. In contrast, the anterior STG remains active throughout the sentence, suggesting a specialization of temporal or salience processing in the posterior STG compared to feature processing in anterior STG (Hamilton et al. 2018). Posterior and anterior auditory cortex has also been suggested to be divided into a dorsal and ventral stream (Bizley and Cohen 2013). In our study, contextual information was present for both when and what sound would follow. A violation of this context activated posterior auditory STG sites, showing that contextual information affects auditory HFB responses differentially following an anterior-posterior division. Combined, these studies and our data point to a specialized role of posterior STG within the auditory stream for implementation of contextual information.

### HFB omission activations are temporally persistent

Another characteristic of the HFB power increases to omissions is that they span several hundred milliseconds in duration. The largest omission HFB amplitudes across subjects are reached at latencies >100ms, peaking between 240ms and 320ms. Omission activations in STG peak ~110ms later compared to auditory activations. For some subjects, significant omission activations occurred in latencies as early as 0ms to 100ms. This was most notable in subject AB01 (Fig. 1B), in which the posterior electrode shows HFB deviating from baseline as early as 0ms to both sounds and omissions.

These omission signals may represent two functional roles with differential temporal profiles: 1) Activations could signify preparatory processes as part of a predictive process, and would be expected to occur early (<100ms); 2) activations could also be an omission response, perhaps signaling surprise in the form of a mismatch or prediction error (Wacongne et al. 2011), and/or auditory saliency detection (Downar et al. 2000). Given our results, we may be seeing both processes in the same region. First, HFB activity may be elevated in the anticipation of a stimulus, which subsequently turns into a surprise signal once the expected stimulus fails to appear. Anticipatory neural firing has been observed in rat auditory cortex in a task manipulating temporal expectations (Jaramillo and Zador 2011). We may be seeing a neural correlate of this rodent finding in HFB in AB01 and IR01, although this preparatory signal was not consistently observed across subjects.

In contrast, HFB activation >100ms to omissions are robust across all subjects with sufficient STG coverage. This activation peaks ~110ms after the auditory-evoked HFB response. This longer latency response may represent a contextual integration process that unfolds over a longer time scale. Such a process may serve to detect saliency including mismatches between what is expected and actual sensory input. Since the posterior STG has been previously implicated with auditory mismatch detection and the ventral attention network (Downar et al. 2001), the HFB omission response is best described as a surprise or mismatch signal.

### Target stimuli elicit increased HFB responses in auditory STG

Target stimuli show larger HFB power increases compared to predictable, non-target stimuli (‘Ba’ and ‘Ga’). Analyses investigating whether such an increase may be larger in auditory-active electrodes that also show omission responses did not show consistent effects. The observed increase likely represents a target detection related attentional modulation (Kastner et al. 1999). It should be noted that contrasting the target ‘Ta’ syllable with non-target ‘Ba’ and ‘Ga’ syllables is not ideal to investigate differences in auditory evoked activity. Syllabic responses have been known to be differentiable in human STG (Chang et al. 2010), and differences could be attributed to differences in the syllabic response in that specific electrode. Moreover, even though the energy in the stimuli were matched, there are differences in spectral properties in the syllable stimuli. Target HFB responses are more reliable in posterior STG/TPJ and inferior frontal cortex in electrodes with no auditory response. The limited coverage of the TPJ and lateral frontal cortex provided some evidence for overlapping omission and target responses in TPJ (Fig. 1B, third plot), and IFG (Fig. 1A, C; Fig. 2B; Supplementary Fig. 2).

### No evidence for stimulus specific information in omission HFB

We found electrodes in posterior STG encoding syllabic information, evident from different amplitudes and time-courses distinguishing between heard syllables (Figs. 1 and 2) and syllable decoding was robust across STG electrodes (Fig. 4). To test if a stimulus template was activated during omissions of expected sounds, we applied a classifier to HFB power time-courses in omission trials, only including electrodes with reliable prediction accuracies for determining which sound was heard. This classification approach was unsuccessful.

HFB activations to omissions were observed only in selective posterior auditory active sites. We therefore re-ran classification of which stimulus was omitted only on electrodes with significant omission HFB increases compared to baseline, including only those time-points within a window of 0-500ms that were delineated in the significant clusters. This classification attempt also remained at chance level, indicating that the omission HFB response in STG does not distinguish between which stimulus was predicted and then omitted.

Based on previous decoding and encoding approaches, the HFB is the most prominent signal containing information on stimulus-specific predictions (Chang et al. 2010; Flinker et al. 2011; Pasley et al. 2012; Martin et al. 2014; Holdgraf et al. 2016). However, it is possible that stimulus-specific expectation information is not carried by activity in the HFB. For example, the gamma (30-70Hz) and beta (15-30Hz) frequency bands have been previously implicated in prediction processes (Arnal and Giraud 2012; Bastos et al. 2012). To test this, we used the power in these two bands to try to decode which stimulus was omitted, which proved unsuccessful. Another explanation could be that such template-specific activations are only occurring in primary auditory cortex which we could not access with our surface electrodes (Kok et al. 2012; Kok et al. 2014). Finally, it is possible that the omission of a sound is more salient than the omission of a *specific* sound. We speculate that the omission response observed here may be related to temporal violations, saliency detection, or both.

### Contextual processing and the posterior STG

A large body of work assigns multitude functions to the posterior STG including speech processing, face processing, audiovisual integration, motion processing and theory of mind (Hein & Knight, 2008). This region is also linked to the ventral attention network, which comprises posterior STG, the temporo-parietal junction (TPJ), inferior frontal gyrus (IFG), insula and cingulate cortex (Downar et al. 2002). This network is linked to identifying salient events, and to re-orientation of attention (Downar et al. 2000; Corbetta and Shulman 2002). This is not surprising, as a deviation from the expectation of what is coming next is core to classifying a stimulus as novel and potentially salient. Contributions from the TPJ and its role in generation of the P300 ERP have been linked to contextual updating (Geng and Vossel 2013). Contextual updating would update the prediction for the next trial, based on the outcome of the current trial. The full cycle of this prediction process follows a time-course that extends beyond early sensory processing. Our data provide insight into the recruitment of auditory regions and their temporal dynamics at different stages of this process. Similar to Downar et al (Downar et al. 2001). we found differential omission activation patterns in the posterior STG/TPJ region and IFG (Figs. 1–3). Some anterior and posterior STG sites were involved specifically in auditory and omission processing, whereas some more posterior sites did not show auditory activations and responded more strongly to targets over omissions (Fig. 1B, third plot). This is in accord with modality-general TPJ activations, whereas the posterior STG is specific to auditory novelty (Downar et al. 2002). The posterior STG may comprise the first node in the network for the detection and response to salient auditory events. In addition, limited sites in IFG were also active to omissions (Fig. 1–3). Given the apparent non-specific, prolonged nature of the HFB omission activation, local neural activity underlying this activation might be involved in binding anticipatory processes with the auditory mismatch processes and the salience detection network.

## Conclusions

We show that omissions of expected sounds elicit a robust HFB increase in a subset of posterior STG auditory active electrodes following a division of anterior vs posterior auditory activity in STG (Ozker et al. 2017; Hamilton et al. 2018). Contrary to current prediction theories, a classification analysis applied to this HFB increase in the STG was unsuccessful in predicting which stimulus was omitted, suggesting that the observed omission HFB activation does not carry stimulus-specific information. Finally, this response is different from that seen in the TPJ, which was shown to respond to both omissions and targets, but not to sounds generally.

## Acknowledgements

Y.M.F and R.T.K. developed the task, interpreted the results and wrote the manuscript. Y.M.F. performed the analyses. A.M. contributed to the classification analyses. R.J. contributed to electrode localization and data visualization. P.B., G.S., and J.L. aided in data acquisition and writing the manuscript. We thank the patients for participating, as well as all people involved in the recording process. We would like to acknowledge all members of the Knight lab for their continued support and feedback, in particular Chris Holdgraf for his MNE Python support, as well as Sandon Griffin for his help with electrode reconstructions. We also thank Kia Nobre for insightful discussions that helped shape the interpretation of our results. We would like to acknowledge our funding sources, including Fulbright and HHMI (Y.M.F.), NIH R37-NS21135 (R.T.K.), P50-MH109429 (R.T.K. and G.S.), P41-EB018783 (G.S.), R01-EB026439 (G.S.), U24-NS109103 (G.S.), U01-NS108916 (G.S.), and R25-HD088157 (G.S.), the US Army Research Office W911NF-15-1-0440 (G.S.), and Fondazione Neurone (G.S.).

## Supplementary figures

**Supplementary Figure 1.**
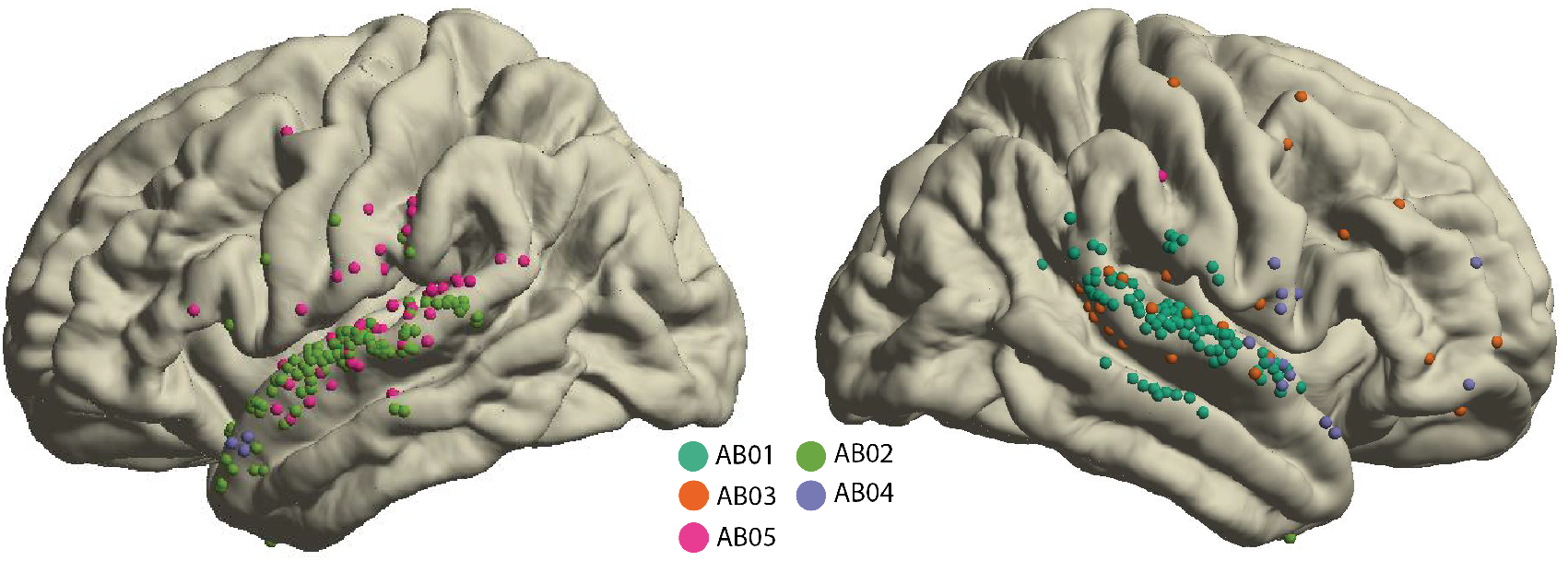
Overview of electrode coverage in all Albany subjects projected onto the fsaverage brain from the Fieldtrip toolbox. Color signifies an individual subject.

**Supplementary Figure 2.**
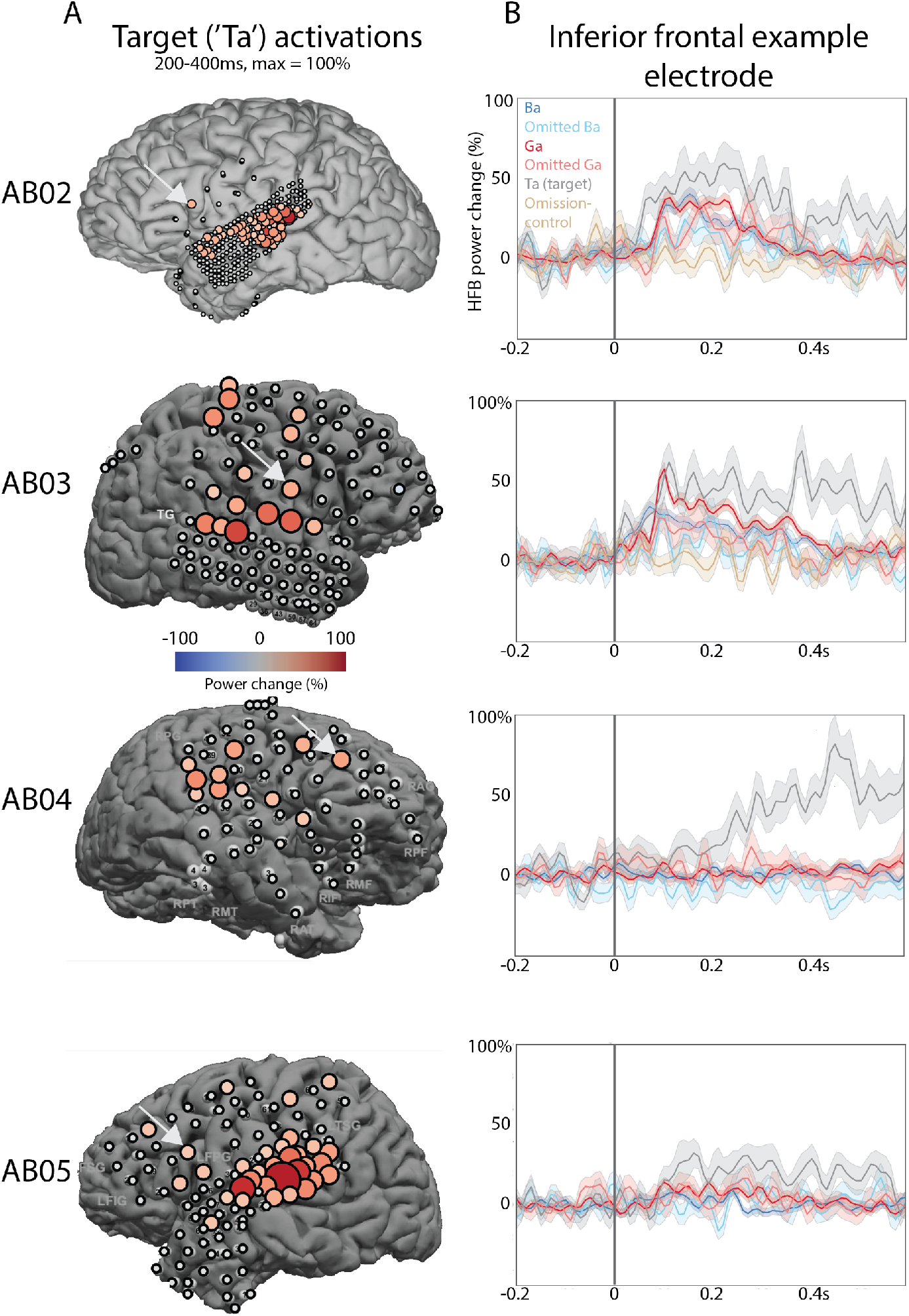
Target HFB power increases in inferior frontal cortex. A) Topography of significant auditory HFB power change. B) Example target-activated electrodes in inferior frontal cortex. Exact location of example electrode is shown by white arrows in topographies in A). Stimulus onset is at 0ms, and traces are HFB responses to ‘Ba’ (dark blue), ‘Ga’ (dark red), omitted ‘Ba’ (light blue), omitted ‘Ga’ (light red), ‘Ta’ (gray) and an omission control (tan).

**Supplementary Figure 3.**
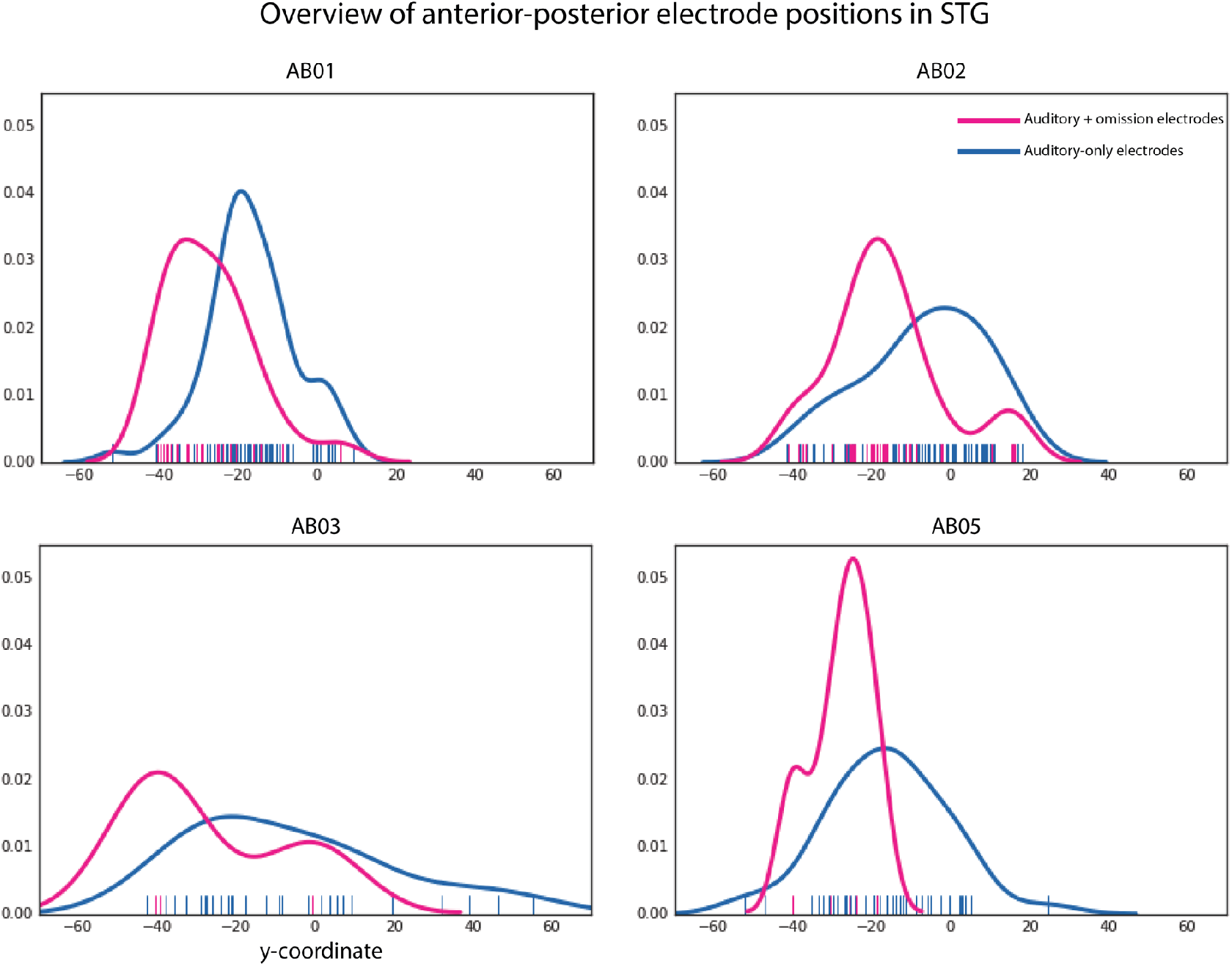
Kernel density plots of the MNI y-coordinates across electrodes for each subject with robust omission activations. This compares the anterior-posterior location of electrodes with both omission as well as auditory HFB activations (magenta) and electrodes with auditory HFB activations but no omission activations (blue). This shows that auditory-active electrodes that are also responsive to omissions consistently are more posterior compared to auditory-active electrodes that do not show omission responses. Colored bars on the y-axis indicate individual electrodes.

**Supplementary Figure 4.**
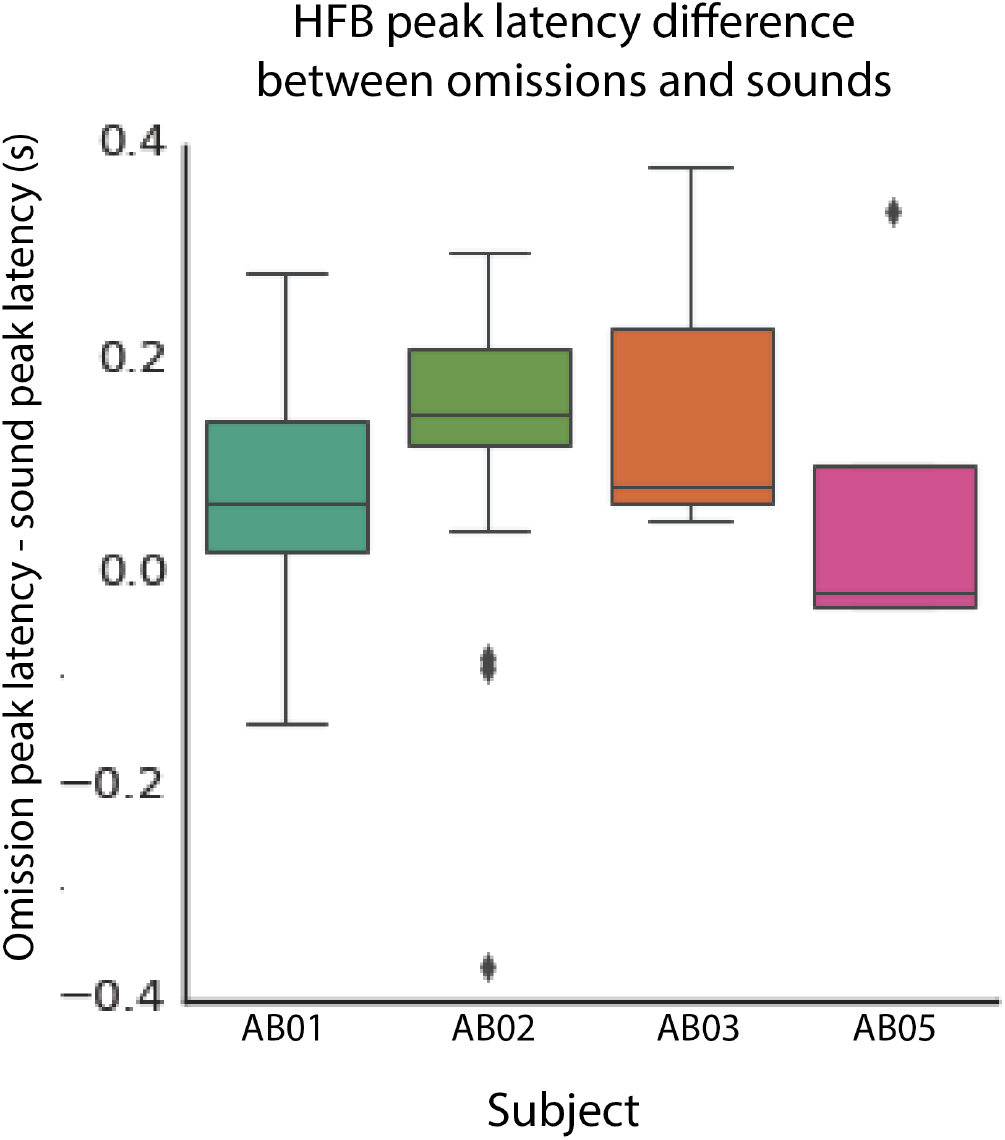
A comparison of peak latencies of the auditory response and omission responses in STG, shown for individual subjects that show both auditory and omission responses in STG. Color codes are the same as in Supplementary Figure 1.

**Supplementary table 1:** Subject demographic information. * denotes unknown

| **Subject** | **Gender** | **Age** | **Handedness** | **Hand used** | **Language lateralization** | **Seizure focus** |
| --- | --- | --- | --- | --- | --- | --- |
| AB01 | F | 33 | Right | Left | * | * |
| AB02 | M | 33 | Right | Right | Left | anterior depth |
| AB03 | M | 57 | Left | Left | Left | * |
| AB04 | M | 69 | Right | Left | * | Anterior ventro-temporal |
| AB05 | M | 31 | Right | Right | * | ventral/temporal pole |
| IR01 | M | 34 | Ambidextrous | Right | * | Bilateral hippocampus |

**Supplementary table 2:** Behavioral data per subject. Reaction times to targets in seconds, hits in percentage, misses and false alarms in number of trials total.

| **Subjects** | **RT median (ms)** | **RT mean (ms)** | **RT standard deviation (ms)** | **Hits** |
| --- | --- | --- | --- | --- |
| AB01 | 643 | 661 | 224 | 98% |
| AB02 | 527 | 563 | 193 | 85% |
| AB03 | 438 | 481 | 160 | 81% |
| AB04 | 550 | 578 | 147 | 92% |
| AB05 | 500 | 528 | 122 | 96% |
| IR01 | 481 | 503 | 91 | 86% |
| **Average** | **523** | **552** | **156** | **90%** |

**Supplementary Table 3:** Prediction accuracies per subject

Omission classification based on electrodes with individual >60% prediction accuracies on 'Ba' vs 'Ga' classification.
| Subjects | Prediction accuracy | Null distribution |  |  |  |
| --- | --- | --- | --- | --- | --- |
|  |  | 50% quantile | 95% quantile | 97.5% quantile | 99.5% quantile |
| AB62 | 50.7% | 50.3% | 60.6% | 62.4% | 65.7% |
| AB81 | 40.4% | 50.2% | 59.6% | 61.4% | 67.6% |
| AB71 | 46.2% | 49.6% | 59.6% | 61.1% | 64.7% |
| AB78 | 44.8% | 49.9% | 60.0% | 62.1% | 64.0% |
| AB79 | 44.3% | 49.9% | 58.6% | 60.8% | 63.2% |
| Mean | 45.28% |  | 59.68% |  |  |
| SD | 0.0371% |  | 0.0073% |  |  |

Omission classification based on significant HFB clusters based on cluster permutation of HFB compared to baseline.
|  | Prediction accuracy | Null distr. |  |  |  |
| --- | --- | --- | --- | --- | --- |
|  |  | 50% quantile | 95% quantile | 97.50% quantile | 99.50% quantile |
| AB01 | 48.8% | 50.4% | 59.0% | 61.0% | 64.5% |
| AB02 | 55.8% | 51.0% | 60.6% | 63.4% | 67.3% |
| AB03 | 38.8% | 50.2% | 60.2% | 61.9% | 65.4% |
| AB04 | 54.8% | 51.4% | 60.2% | 62.0% | 65.2% |
| AB05 | 46.5% | 49.5% | 58.1% | 61.0% | 64.6% |
| Mean | 48.9% |  | 59.6% |  |  |
| SD | 0.0689% |  | 0.0104% |  |  |

Prediction accuracies of heard 'Ba' vs 'Ga' based on significant HFB activations
|  | Prediction<br>Accuracy | Null distr. |  |  |  |
| --- | --- | --- | --- | --- | --- |
|  |  | 50%<br>quantile | 95%<br>quantile | 97.50%<br>quantile | 99.50%<br>quantile |
| AB01 | 96.0% | 49.9% | 53.5% | 53.8% | 55.0% |
| AB02 | 89.3% | 50.2% | 53.5% | 54.0% | 55.2% |
| AB03 | 73.4% | 49.9% | 53.0% | 53.8% | 54.9% |
| AB04 | 66.7% | 50.0% | 52.5% | 53.2% | 54.5% |
| AB05 | 96.6% | 49.9% | 53.1% | 53.8% | 54.8% |
| Mean | 84.4% |  |  |  | 54.9% |
| SD | 0.1362% |  |  |  | 0.0026% |

